# Changes in neuronal immunofluorescence in the C- versus N-terminal domains of hnRNP H following D1 dopamine receptor activation

**DOI:** 10.1101/305565

**Authors:** Qiu T. Ruan, Neema Yazdani, Jacob A. Beierle, Kathryn M. Hixson, Kristen E. Hokenson, Daniel J. Apicco, Kimberly P. Luttik, Karen Zheng, Brandon F. Maziuk, Peter E. A. Ash, Karen K. Szumlinski, Shelley J. Russek, Benjamin Wolozin, Camron D. Bryant

**Affiliations:** Laboratory of Addiction Genetics, Department of Pharmacology and Experimental Therapeutics and Psychiatry, Boston University School of Medicine; Biomolecular Pharmacology Training Program, Department of Pharmacology and Experimental Therapeutics, Boston University School of Medicine; Transformative Training Program in Addiction Science, Boston University; Laboratory of Translational Epilepsy, Department of Pharmacology and Experimental Therapeutics and Biology, Boston University School of Medicine; Laboratory of Neurodegeneration, Department of Pharmacology and Experimental Therapeutics and Neurology, Boston University School of Medicine; Department of Psychological and Brain Sciences, University of California, Santa Barbara

**Author notes:** These authors contributed equally to the work (Qiu T. Ruan, Neema Yazdani). Corresponding Author, 72 E. Concord Street, L-606C, Boston, MA 02118.

**Keywords:** hnRNP, RNA binding protein, psychostimulant use disorder, amphetamine, cocaine, addiction

## Abstract

RNA binding proteins are a diverse class of proteins that regulate all aspects of RNA metabolism. Accumulating studies indicate that heterogeneous nuclear ribonucleoproteins are associated with cellular adaptations in response to drugs of abuse. We recently mapped and validated heterogeneous nuclear ribonucleoprotein H1 (*Hnrnph1*) as a quantitative trait gene underlying differential behavioral sensitivity to methamphetamine. The molecular mechanisms by which hnRNP H1 alters methamphetamine behaviors are unknown but could involve preand/or post-synaptic changes in protein localization and function. Methamphetamine initiates post-synaptic D1 dopamine receptor signaling indirectly by binding to pre-synaptic dopamine transporters and vesicular monoamine transporters of midbrain dopaminergic neurons which triggers revers e transport and accumulation of dopamine at the synapse. Here, we examined changes in neuronal localization of hnRNP H in primary rat cortical neurons that express dopamine receptors that can be modulated by the D1 or D2 dopamine receptor agonists SKF38393 and (-)-Quinpirole HCl, respectively. Basal immunostaining of hnRNP H was localized primarily to the nucleus. D1 dopamine receptor activation induced an increase in hnRNP H nuclear immunostaining as detected by immunocytochemistry with a C-domain directed antibody containing epitope near the glycine-rich domain but not with an N-domain specific antibody. Although there was no change in hnRNP H protein in the nucleus or cytoplasm, there was a decrease in *Hnrnph1* transcript following D1 receptor stimulation. Taken together, these results suggest that D1 receptor activation increases availability of the hnRNP H C-terminal epitope, which could potentially reflect changes in protein-protein interactions. Thus, D1 receptor signaling could represent a key molecular post-synaptic event linking *Hnrnph1* polymorphisms to drug-induced behavior.

**Highlights:**

- We investigated immunofluorescence and localization of hnRNP H following dopamine receptor activation in primary rat cortical neurons.
- Activation of D1 dopamine receptor increased nuclear immunofluorescence of the C-terminal domain but not the N-terminal domain of hnRNP H with no change in either nuclear or cytoplasmic levels of hnRNP H protein.
- D1 dopamine signaling could alter interaction of hnRNP H with other proteins.

## 1. Introduction

Heterogeneous nuclear ribonucleoproteins (hnRNPs) represent a large family of RNA binding proteins (RBPs) with multifunctional roles in RNA biogenesis and metabolism, both in the nucleus and the cytoplasm [1,2]. In the nucleus, hnRNPs regulate transcription, splicing, and mRNA stability [1–4]. In the cytoplasm, hnRNPs can repress or enhance translation by binding to 3’- and 5’-untranslated regions of mRNAs [1,2,5]. Several hnRNPs are present in both the nucleus and cytoplasm and bind to mRNA targets to regulate bidirectional transport [6,7]. In response to neuronal stimulation, hnRNPs can also regulate synaptic plasticity through shuttling mRNAs encoding synaptodendritic proteins, kinases, and cytoskeletal elements to dendritic processes [4,8–10].

The hnRNP H family of RBPs including, H1, H2, and F, regulate mRNA alternative splicing and polyadenylation [3,12–14]. hnRNP H1 and H2 share a 96% protein sequence homology and have yet to be functionally differentiated [6]. hnRNP H contains three quasi-RNA recognition motifs (qRRMs), and two glycine-rich domains, GY and GYR [7,15]. The qRRMs recognize and bind to poly-G runs of varying lengths or GGG triplets within RNA sequences [15,16]. The glycine-rich GYR domain is critical for the nuclear localization of hnRNP H [7], while both GYR and GY domains facilitate protein-protein interactions [5].

We used quantitative trait locus (QTL) mapping, positional cloning, and gene editing to identify *Hnrnph1* as a quantitative trait gene underlying reduced sensitivity to the locomotor stimulant response to methamphetamine (MA) in mice [17]. Based on transcriptome analysis of the QTL within the striatum, we hypothesized that the mechanism could involve a deficit in mesolimbic dopaminergic neuron development. Dopaminergic neurons in the ventral midbrain project to the basal forebrain and prefrontal cortex where MA binds to pre-synaptic dopamine transporters and vesicular monoamine transporters and induces reverse transport, increased synaptic dopamine, and activation of synaptic D1 Gs and D2 Gi/Go-coupled receptors [18,19]. hnRNP H could potentially act in presynaptic dopaminergic neurons or in postsynaptic forebrain neurons to influence the acute behavioral response to MA and activity-dependent neurobehavioral plasticity. In the present study, we utilized primary rat cortical neurons as an *in vitro* model to examine changes in localization of hnRNP H following dopamine receptor activation.

## 2. Materials and Methods

### 2.1. Primary rat cortical neuron culture and neuronal treatment

Primary rat cortical neurons were dissected from neocortex of E18 Sprague-Dawley embryos (Charles River Laboratories, Wilmington, MA, USA) and grown in media as described [20]. Dissociated neurons were plated on Poly-L-lysine coated dishes (18k cells/cm^2^) and placed in an incubator (37ºC; 5% CO_2_) for attachment to coverslips/dish surface. Neuronal cultures were grown for 8 days *in vitro* (DIV8) until treatment unless specified otherwise (DIV16). For control treatment, 1 ml of conditioned media was replaced with 1 ml of warmed neurobasal media. For experimental treatments, 1 ml warmed neurobasal media was mixed with potassium chloride (KCl; Boston BioProducts, Ashland, MA, USA), the D1 dopamine receptor agonist SKF38393 (Sigma Aldrich, St. Louis, MO, USA), the D1 dopamine receptor antagonist SCH23390 (Sigma Aldrich) plus SKF38393 or the D2 dopamine receptor agonist (-)-Quinpirole HCl (Quinpirole; Sigma Aldrich). Final treatment concentrations were 1 µM SKF38393, 1 µM Quinpirole and 10 nM SCH23390 and cells were incubated for 1.5 h. Refer to Supplement for detailed description of immunocytochemistry protocol and quantification of immunofluorescence. To test for the specificity of the hnRNP H antibody, immunoabsorption using a custom-made blocking peptide to the C-term of the antibody was performed (see Supplement for additional details).

### 2.2. Nuclear/cytoplasmic fractionation and immunoblotting

Following D1 dopamine receptor stimulation with SKF38393, neurons were fractionated using Pierce^TM^ NE-PER Nuclear and Cytoplasmic Extraction Reagents (Thermo Scientific, Waltham, MA, USA). Fractionation efficiency was assessed through immunologically probing for CREB (a nuclear marker) and HSP90 (a cytoplasmic marker). Following densitometry analysis (using ImageJ), the quantified values for the hnRNP H bands were normalized to their respective loading control bands (CREB for nucleus fractions and HSP90 for cytoplasmic fractions) to generate final comparable ‘arbitrary unit’ (AU) protein level values. AU values were averaged and normalized to the control group to calculate fold-change. Two-way ANOVA was used to assess interaction between cellular compartment and treatment.

## 3. Results

### 3.1. Increase in hnRNP H nuclear immunofluorescence at the C-terminus but not the N-terminus following D1 dopamine receptor activation

We previously reported an increase in nuclear fluorescence of hnRNP H following 2 h of KCl stimulation using ICC with an antibody against the hnRNP H C-domain [21]. We replicated this result following 1.5 h of KCl stimulation via ICC against the hnRNP H C-domain (GY; **Fig. S1A, B**). Strikingly however, KCl stimulation followed by ICC against the hnRNP H N-domain (qRRM 1-2; **Fig. S2A**) revealed no change in nuclear fluorescence of hnRNP H (**Fig. S1C**). Taken together, our results suggest that the differential detection of hnRNP H by these two antibodies at the C versus N terminus could reflect activity-dependent changes in hnRNP H function.

We next asked whether dopamine receptor activation would induce an increase in hnRNP H fluorescence by treating neurons with D1 or D2 dopamine receptor agonists. Similar to depolarization with KCl, neurons treated with the D1 dopamine receptor agonist SKF38393 showed an increase in nuclear fluorescence of hnRNP H at the C-terminus. Furthermore, this increase was reversed by co-treatment with the D1 dopamine receptor antagonist SCH23390 (**Fig. 1B, C**). In contrast, there was no increase in nuclear immunofluorescence with the N-domain antibody following D1 dopamine receptor stimulation (**Fig. 1D**). Neurons treated with the D2 receptor agonist, Quinpirole-HCl, did not show any change in hnRNP H nuclear immunofluorescence with the C-domain antibody (**Fig. S2**), although this null result could potentially be explained by lower D2 versus D1 receptor expression. In support, *Drd2* transcript abundance was lower than *Drd1* in DIV8 primary rat cortical neurons (**Fig. S3A**). D1 dopamine receptor activation did not significantly affect transcription of *Drd1* or *Drd2* (**Fig. S3B**). In primary rat cortical neurons, we detected D1 dopamine receptor immunoreactivity (**Fig. S3C**). To determine whether the D1-mediated increase in C-terminal staining of hnRNP H depended on the age of the neurons, we also treated DIV16 primary rat cortical neurons with SKF38393 that also increased C-terminal hnRNP H immunofluorescence (**Fig. S4)**.

**Fig. 1.**
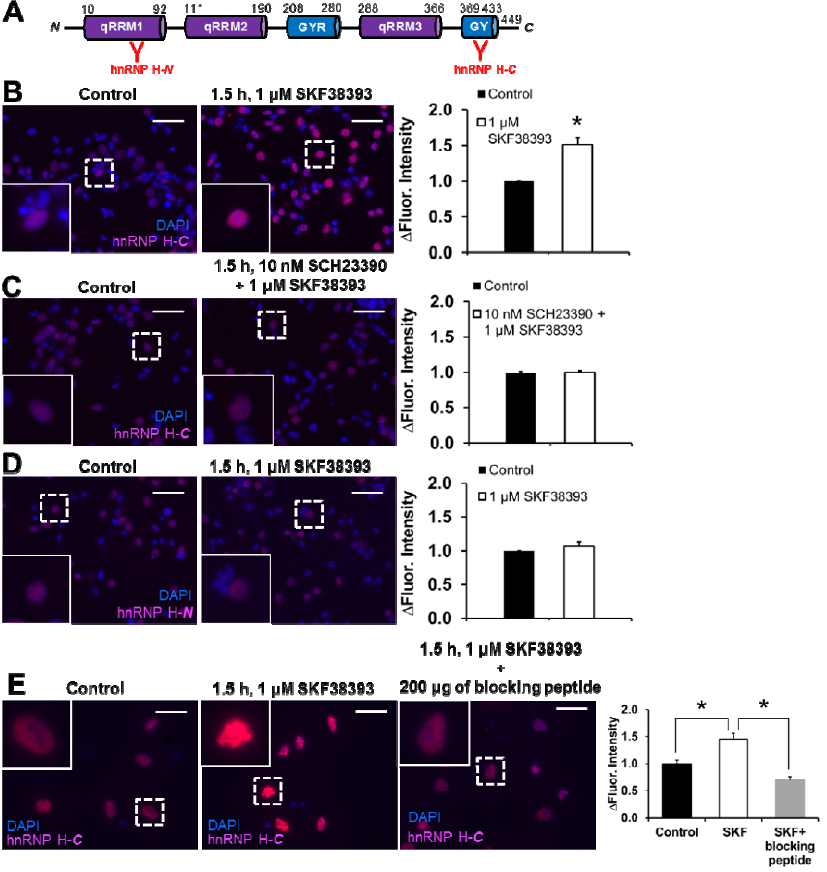
Increase in hnRNP H nuclear immunofluorescence at the C-terminus but not the N-terminus following D1 dopamine receptor activation. DIV8 primary rat cortical neurons were treated with vehicle (Control, n = 4), 1 μM of SKF38393 (D1 dopamine receptor agonist, n = 4) and/or 10 nM of SCH23390 (D1 dopamine receptor antagonist n = 5) for 1.5 h. **(A)** Schematic shows hnRNP H and its quasi-RNA recognition motifs (qRRMs) and glycine-rich domains (GYR and GY). The red “Y” markings indicate the relative antibody epitopes in the N-domain or C-domain regions of hnRNP H. **(B,C)** ICC and intensity analysis revealed a significant increase in C-domain staining of nuclear hnRNP H following 1 μM SKF38393 [unpaired Student’s t-test, t_6_ = −4.94, *\**p = 0.003] that was blocked by co-administration of 1 μM SKF38393 and 10 nM SCH23390 [unpaired Student’s t-test, t_7_ = −0.34, p = 0.74]. **(D)** No significant difference was found in N-domain immunoreactivity of nuclear hnRNP H following 1 μM SKF38393 treatment [unpaired Student’s t-test, t_6_ = −1.12, p = 0.30]. **(E)** There was an effect of treatment on C-domain staining of hnRNP H [One-way ANOVA, F_2,6_ = 20.38, p < 0.001, n = 3 per condition]. SKF38393-induced increase in hnRNP H nuclear immunofluorescence was reduced in the present of the blocking peptide [post-hoc Bonferroni correction for Control vs SKF+blocking peptide: t_4_ = 5.849; *p = 0.004]. A significant increase in C-domain staining of nuclear hnRNP H following 1 μM SKF38393 [post-hoc Bonferroni correction for Control vs SKF: t_4_ = −3.454; *p = 0.025]. Black bar = Control; white bar = SKF treatment. Images were obtained on a Zeiss AxioObserver microscope with the 20X objective under uniform settings. Scale bars represent 40 µm.

Our findings were dependent on an hnRNP H antibody that recognizes the epitope at the C-terminus. This epitope is located in a region between residues 400 to 488 of the human hnRNP H1 as indicated by the numbering provided in entry NP_005511.1 (Bethyl Laboratories, Montgomery, TX, USA). To provide further evidence that this region was responsible for the D1-mediated increase in C-terminal staining of hnRNP H, we conducted immunoabsorption with a blocking peptide that binds to the C-terminal end of the hnRNP H protein prior to primary antibody incubation for ICC. As expected, there was a complete blockade of D1-induced increase in nuclear immunofluorescence of hnRNP H (**Fig. 1E**), which is consistent with increased hnRNP H epitope availability following D1 dopamine receptor stimulation.

Using co-ICC of the neuron-specific marker NeuN (e.g. RBFOX3), hnRNP H and DAPI, we determined that hnRNP H immunofluorescence was restricted to the nucleus and localized to neurons (**Figs. 2A-B & S5**). An increase in D1 dopamine receptor agonist-induced increase in C-domain hnRNP H nuclear immunofluorescence was detected in NeuN+/hnRNP H+ neurons (**Fig. 2A-B**). In addition to neuronal cells (60%), we also identified a population of DAPI-stained nuclei in both control and treated (1 uM SKF38393) that did not colocalize with hnRNP H or NeuN (40%) (**Fig. 2C & S5**). NeuN is used as a marker for differentiated, post-mitotic neurons but in contrast to nestin, NeuN is not expressed in undifferentiated neuronal precursors [22]. Thus, we are underestimating the total number of neurons that we previously estimated to be 80% using nestin in this neuronal culture system by the Russek lab. Importantly, there was no difference in the number of NeuN/hnRNP H-positive or non-NeuN staining cells between treatments (**Figs. 2C & S5**).

**Fig. 2.**
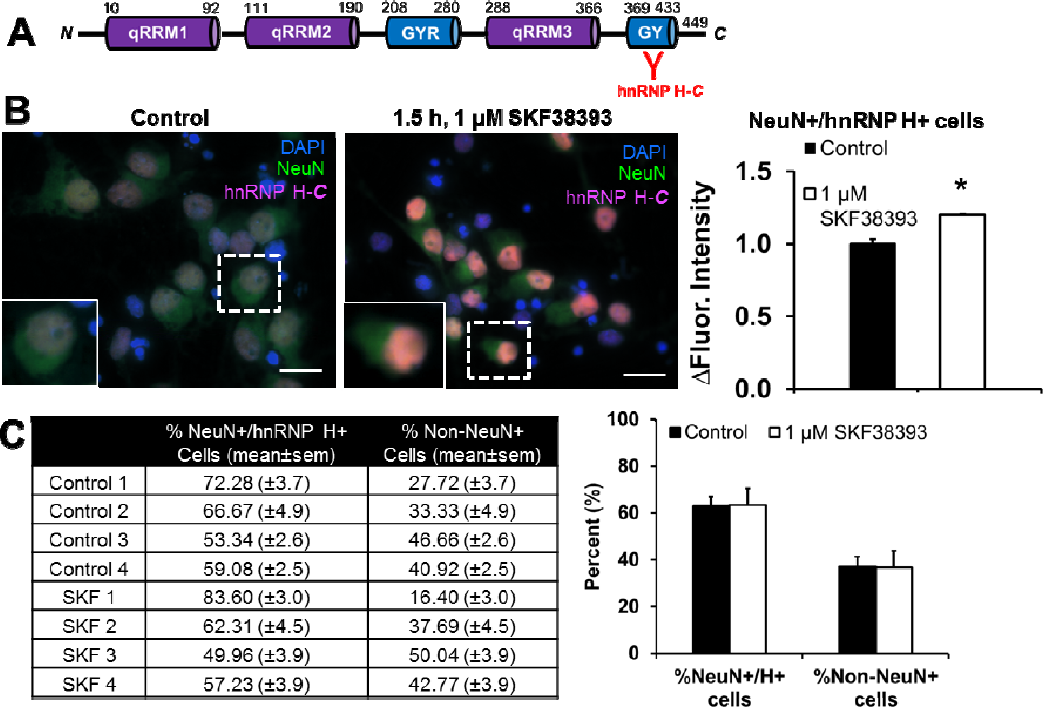
Increase in C-domain hnRNP H nuclear immunofluorescence of NeuN-positive neurons. DIV8 primary rat cortical neurons were treated with media (Control, n = 4) or 1 μM of SKF38393 (D1 dopamine receptor agonist, n = 4) for 1.5 h. **(A)** Schematic shows hnRNP H domains and the red “Y” marking indicates the relative antibody epitopes in the C-domain regions of hnRNP H. **(B)** Merged images showing co-localization of hnRNP H with NeuN (neuronal marker) and DAPI (nuclear marker). Images from individual channel are shown in Fig S5. hnRNP H staining was localized to the nucleus and no cytoplasmic localization of hnRNP H was detected. ICC and intensity analysis revealed a significant increase in C-domain staining of nuclear hnRNP H following 1 μM SKF38393 for NeuN-positive and hnRNP H-positive neurons (unpaired Student’s t-test, t_6_ = −5.58, *p < 0.01). **(C)** There was co-localized staining of hnRNP H with NeuN-positive neuronal staining and no detectable staining of hnRNP H staining in non-neuronal cells. No significant differences in percentage of NeuN/hnRNP H-positive cells [unpaired Student’s t-test, t_6_ = −0.052, p = 0.96] or non-NeuN cells [unpaired Student’s t-test, t_6_ = −0.052, p = 0.96] were noted between treatment conditions. Black bar = Control; white bar = SKF treatment. Images were obtained on a Zeiss AxioObserver microscope with the 63X objective under uniform settings. Scale bars represent 15 µm.

### 3.2. Changes in *Hnrnph1* and *Hnrnph2* transcription or hnRNP H protein expression following D1 dopamine receptor stimulation

D1 receptor activation induced a small but significant decrease in *Hnrnph1* mRNA levels but no change in *Hnrnph2* mRNA (**Fig. 3A**). KCl treatment also had no effect on transcription of either *Hnrnph1* or *Hnrnph2* (**Fig. S6A**). There was also no change in transcription of other components of the hnRNP H family [23] including *Hnrnph3*, *Hnrnpf*, and *Grsf1* **(Fig. S8**). Interestingly, although not detectable via ICC, immunoblot revealed the presence of a small amount of hnRNP H protein in the cytoplasmic fraction compared to the nuclear fraction (**Figs. 3B & S6B**). However, there was no change in hnRNP H protein in the nucleus or cytoplasm following D1 dopamine receptor stimulation (**Fig. 3B**). Importantly, neither the C- nor the N-domain directed antibody displayed any non-specific binding and banding at baseline or in response to SKF38393 treatment (**Fig. S7A, B**), therefore further supporting specificity of hnRNP H detection. In addition, immunoprecipitation of C-term followed by immunoblotting with the N-term hnRNP H antibody demonstrated that the two antibodies are capable of recognizing the same protein (**Fig. S7C**).

**Fig. 3.**
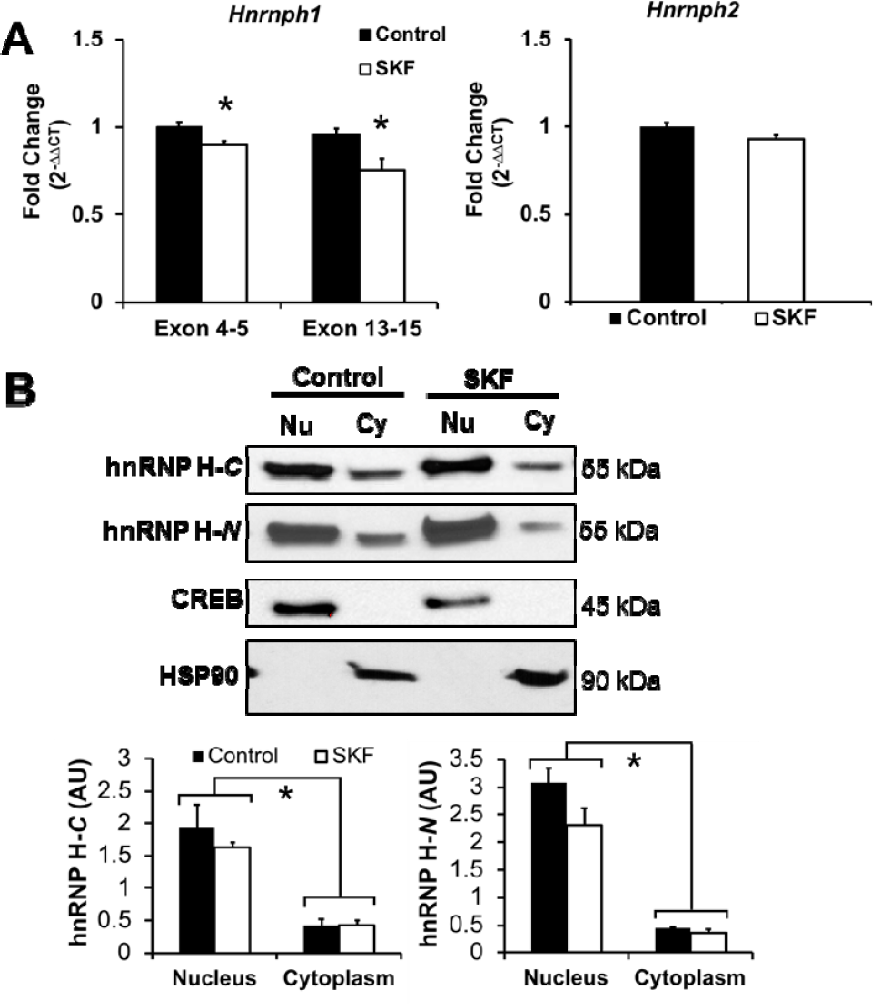
Changes in *Hnrnph1* and *Hnrnph2* transcript abundance or in hnRNP H protein in response to D1 dopamine receptor activation. DIV8 primary rat cortical neurons were treated with vehicle (Control, n = 4) or 1 μM of SKF38393 (D1 dopamine receptor agonist, n = 4) for 1.5 h. **(A)** qPCR analysis revealed a small decrease in *Hnrnph1* mRNA (*top left*) [Exon 4-5: unpaired Student’s t-test, t_6_ = 2.866, *\**p = 0.03; Exon13-15: unpaired Student’s t-test, t_6_ = 3.367, *p=0.02] or *Hnrnph2* mRNA (*top right*) [unpaired Student’s t-test, t_6_ = 2.432, p = 0.051]. **(B)** Immunoblot analysis of fractionated samples indicated no significant change in hnRNP H protein levels in the nucleus (Nu) or cytoplasm (Cy), as detected by either C-domain (*left*) or N-domain (*right*) antibodies. Two-way ANOVA with Cellular Compartment and Treatment as factors revealed a significant effect of Cellular Compartment [C-domain: F(1,12) = 51.45, p = 1.13 × 10^−5^; N-domain: F(1,12) = 123.52, p = 1.13 × 10^−7^], no significant effect of Treatment [C-domain: F(1,12) = 1.06, p = 0.32; N-domain: F(1,12) = 4.11, p = 0.07], and no interaction [C-domain: F(1,12) = 0.26, p = 0.62; N-domain: F(1,12) = 2.66, p = 0.13]. Successful nuclear and cytoplasmic fractionation was confirmed by CREB and HSP90. For either C- and N-domain immunoblot study: Control n = 4, 1 μM SKF38393: n = 4. Black bar = Control; white bar = SKF treatment.

### 3.3. Blue native gel analysis of changes in hnRNP H-associated protein complexes following D1 dopamine receptor activation in rat cortical primary neurons

To test whether changes in protein complexes containing hnRNP H were associated with the D1 dopamine receptor-induced C-domain-specific increase in immunofluorescence staining in hnRNP H and thus, whether this could potentially explain the increase in epitope availability, we conducted native gel electrophoresis and immunoblotted with the hnRNP H (C-term) antibody. There were no visible changes in the banding pattern or intensity of the bands representing hnRNP H complexes (**Fig. S9**). The current results do not support a change in hnRNP H protein complexes in response to D1 receptor activation that could account for the increase in epitope availability at the C-terminus of hnRNP H.

## 4. Discussion

Neuronal depolarization with KCl ([21] & **Fig. S1**) and D1 dopamine receptor activation induced a robust increase in nuclear fluorescence of hnRNP H in primary rat cortical neurons (**Fig. 1**). Previous studies demonstrated that KCl stimulation of neurons induced the translocation and accumulation of RNA-binding proteins (RBPs) such as HuD and hnRNP A2 from the nucleus to dendritic processes [10,24]. Although several hnRNPs exhibit activity-dependent neuronal translocation, we did not detect any gross change in hnRNP H localization from the nucleus to the cytoplasm in response to neuronal depolarization or dopamine receptor activation (**Figs. S1 & 1**). Interestingly, we detected a low level of hnRNP H in the cytoplasmic fraction of DIV8 primary rat cortical neurons (**Figs. S6 & 3**). Cytoplasmic hnRNP H was reported using myc-tagged hnRNP H in COS7 cells [25]. Here, we extended this observation to rat cortical neurons and thus, both nuclear (e.g., splicing) and cytoplasmic functions of hnRNP H (e.g., translational regulation) could contribute to activity-dependent synaptic plasticity.

An important question is whether the SKF38393-mediated increase in nuclear fluorescence of hnRNP H finding is specific to the DIV of primary rat cortical neurons. hnRNP H transcript and protein levels in primary rat cortical neurons have been shown to decrease between DIV 1 through 15, which correlated negatively with neuronal differentiation [26]. The decrease in hnRNP H over time could potentially explain the less pronounced but still significant effect of SKF38393 treatment on the nuclear fluorescence of hnRNP H in DIV16 neurons (**Fig. S4**). Thus, the nuclear hnRNP H immunofluorescence following SKF38393 and KCl treatments could vary with neuronal age and maturation.

The C-domain of hnRNP H contains glycine-rich domains (GYR and GY; **Fig. 1A**) which mediate protein-protein interactions [7]. Both D1 receptor and KCl stimulation induced an increase in nuclear immunofluorescence of hnRNP H, as detected by an antibody that binds to the GY domain of the C-terminus (**Fig. 1B; Fig. S1B**). This increased fluorescence occurred without any change in protein level (**Fig. 3; Fig. S6**). We used BN PAGE to test whether increased immunofluorescence was associated with a change in hnRNP H-associated protein complexes but did not observe any visible changes (**Fig. S9**). The similarly-sized protein complexes in treated versus untreated neurons could be heterogeneous and differ in weight and composition at a higher resolution of analysis. Future *in vivo* studies with D1 agonists combined with immunoprecipitation and mass spectrometry will determine whether hnRNP H protein complexes change in molecular weight and composition.

It is possible that depolarization or D1 receptor activation disrupted ribonucleoprotein complexes containing hnRNP H, perhaps through a post-translational modification that increased epitope availability in the C-domain. Interestingly, both KCl-mediated neuronal depolarization and D1 receptor stimulation activate adenylate cyclase activity and increase the level of cytoplasmic cyclic adenosine monophosphate (cAMP), leading to activation of several kinases, including protein kinase A (PKA) [27,28]. In maintaining energy homeostasis, activation of PKA results in its translocation to the nucleus where it can phosphorylate hnRNP H, leading to increased glucose uptake [29]. There are several predicted phosphorylation sites of hnRNP H [30]. Phosphorylation of hnRNP A1, I, K and L can alter RBP nucleotide-binding capacity, nucleocytoplasmic shuttling, ribonucleoprotein complexes, and/or alternative splicing [6,31–35]. Notably, modifications such as mutations in the GYR domain nuclear-localization sequence of hnRNP H2 could cause cytoplasmic retention and perturb neurodevelopment in females [7,36]. Mutations in the glycine-rich domains of other RBPs, such as FUS, TDP-43, hnRNPA1 and A2B1 could induce aberrant ribonucleoprotein complexes, contributing to neurodegenerative disease [37]. Future phosphoproteomic studies examining phosphorylation states at the glycine-rich C-terminus of hnRNP H following D1 dopamine receptor activation will identify phosphorylation sites that can be studied experimentally with the development of phospho-specific hnRNP H antibodies. These findings could inform the mechanisms underlying the observed D1-induced increase in C-domain epitope availability and hypothesized changes in protein-protein interactions.

We previously proposed that hnRNP H could regulate mesocorticolimbic dopaminergic neuron development and function [17,21]. Alcohol-binge drinking in mice identified a correlation between alcohol intake and *Hnrnph1*, *Hnrnpm*, and *Hnrnpc* expression in dopaminergic neurons of the ventral tegmental area, implicating a potential role of hnRNPs in dopaminergic neuroadaptive changes [38]. In the context of psychostimulant addiction, we recently identified *Hnrnph1* as a quantitative trait gene underlying differential sensitivity to MA [17]. It is unclear whether hnRNP H1 functions at a presynaptic or postsynaptic level to modulate behavioral responses to drugs of abuse. Our current observations raise the possibility that post-synaptic D1 receptor signaling could initiate acute and chronic downstream cellular adaptations via hnRNP H. Our findings warrant additional proteomic studies to clarify the dynamics of pre-synaptic versus post-synaptic hnRNP H-associated protein complexes *in vivo* during D1 dopamine receptor signaling.

## 5. Conclusion

We observed a selective, D1 dopamine receptor-dependent increase in nuclear immunofluorescence of the C-domain, but not the N-domain, of hnRNP H without changes in nuclear or cytoplasmic levels of hnRNP H protein. D1-dependent dopamine signaling could affect the interaction of hnRNP H with other proteins and mRNAs and contribute to long-term neurobehavioral adaptations in response to psychostimulants. Our findings warrant further investigation into potential changes in protein-protein interactions with hnRNP H in response to dopamine receptor stimulation.

## Acknowledgments

This study was supported by R01DA039168 (C.D.B.), F31DA40324 (N.Y.), T32GM008541 (N.Y., Q.T.R., K.E.H., K.M.H., D.J.A., J.A.B.), Burroughs Wellcome Fund Transformative Training Program in Addiction Science #1011479 (N.Y., Q.T.R., J.A.B.), F31NS106751 (B.F.M), R01NS051710 (S.J.R.), R21NS083057 (S.J.R.), and R01AG050471 (B.W.).

## References

[1] S.P. Han, Y.H. Tang, R. Smith, Functional diversity of the hnRNPs: past, present and perspectives., Biochem. J. 430 (2010) 379–392. doi:10.1042/BJ20100396.

[2] T. Geuens, D. Bouhy, V. Timmerman, The hnRNP family: insights into their role in health and disease, Hum. Genet. 135 (2016) 851–867. doi:10.1007/s00439-016-1683-5.

[3] G.K. Arhin, M. Boots, P.S. Bagga, C. Milcarek, J. Wilusz, Downstream sequence elements with different affinities for the hnRNP H/H′ protein influence the processing efficiency of mammalian polyadenylation signals, Nucleic Acids Res. 30 (2002) 1842–1850.

[4] G. Zhang, T.A. Neubert, B.A. Jordan, RNA binding proteins accumulate at the postsynaptic density with synaptic activity, J. Neurosci. 32 (2012) 599–609. doi:10.1523/JNEUROSCI.2463-11.2012 [doi].

[5] P.J. Uren, E. Bahrami-Samani, P.R. de Araujo, C. Vogel, M. Qiao, S.C. Burns, A.D. Smith, L.O.F. Penalva, High-throughput analyses of hnRNP H1 dissects its multi-functional aspect., RNA Biol. 0 (2016) 1–12. doi:10.1080/15476286.2015.1138030.

[6] B. Honoré, H.H. Rasmussen, H. Vorum, K. Dejgaard, X. Liu, P. Gromov, P. Madsen, B. Gesser, N. Tommerup, J.E. Celis, Heterogeneous nuclear ribonucleoproteins H, H′, and F are members of a ubiquitously expressed subfamily of related but distinct proteins encoded by genes mapping to different chromosomes. J Biol. Chem. 270 (1995) 28780–28789. doi:10.1074/jbc.270.48.28780.

[7] C.M. Van Dusen, L. Yee, L.M. McNally, M.T. McNally, A glycine-rich domain of hnRNP H/F promotes nucleocytoplasmic shuttling and nuclear import through an interaction with transportin 1., Mol. Cell. Biol. 30 (2010) 2552–2562. doi:10.1128/MCB.00230-09.

[8] J.C. Darnell, Defects in translational regulation contributing to human cognitive and behavioral disease, Curr. Opin. Genet. Dev. 21 (2011) 465–473. doi:10.1016/j.gde.2011.05.002.

[9] S.Y. Grooms, K.-M. Noh, R. Regis, G.J. Bassell, M.K. Bryan, R.C. Carroll, R.S. Zukin, Activity bidirectionally regulates AMPA receptor mRNA abundance in dendrites of hippocampal neurons, J. Neurosci. v26 (2006) 8339–8351. doi:10.1523/JNEUROSCI.0472-06.2006.

[10] I.A. Muslimov, A. Tuzhilin, T.H. Tang, R.K.S. Wong, R. Bianchi, H. Tiedge, Interactions of noncanonical motifs with hnRNP A2 promote activity-dependent RNA transport in neurons, J. Cell Biol. 205 (2014) 493–510. doi:10.1083/jcb.201310045.

[11] G. Dreyfuss, M.J. Matunis, S. Pinol-Roma, C.G. Burd, hnRNP Proteins and the Biogenesis of mRNA, Annu. Rev. Biochem. 62 (1993) 289–321. doi:10.1146/annurev.biochem.62.1.289.

[12] S.C. Huelga, A.Q. Vu, J.D. Arnold, T.D. Liang, P.P. Liu, B.Y. Yan, J.P. Donohue, L. Shiue, S. Hoon, S. Brenner, M. Ares, G.W. Yeo, Integrative Genome-wide Analysis Reveals Cooperative Regulation of Alternative Splicing by hnRNP Proteins, Cell Rep. 1 (2012) 167–178. doi:10.1016/j.celrep.2012.02.001.

[13] Y. Katz, E.T. Wang, E.M. Airoldi, C.B. Burge, Analysis and design of RNA sequencing experiments for identifying isoform regulation, Nat. Methods. 7 (2010) 1009–1015. doi:10.1038/nmeth.1528.

[14] E. Wang, V. Aslanzadeh, F. Papa, H. Zhu, P. de la Grange, F. Cambi, Global Profiling of Alternative Splicing Events and Gene Expression Regulated by hnRNPH/F, PLoS One. 7 (2012) e52166. doi:10.1371/journal.pone.0051266.1

[15] C. Dominguez, F.H.T. Allain, NMR structure of the three quasi RNA recognition motifs (qRRMs) of human hnRNP F and interaction studies with Bcl-x G-tract RNA: A novel mode of RNA recognition, Nucleic Acids Res. (2006). doi:10.1093/nar/gkl488.

[16] X. Xiao, Z. Wang, M. Jang, R. Nutiu, E.T. Wang, C.B. Burge, Splice site strength–dependent activity and genetic buffering by poly-G runs, Nat. Struct. Mol. Biol. 16 (2009) 1094–1100. doi:10.1038/nsmb.1661.

[17] N. Yazdani, C.C. Parker, Y. Shen, E.R. Reed, M.A. Guido, L.A. Kole, S.L. Kirkpatrick, J.E. Lim, G. Sokoloff, R. Cheng, W.E. Johnson, A.A. Palmer, C.D. Bryant, Hnrnph1 Is A Quantitative Trait Gene for Methamphetamine Sensitivity, PLoS Genet. 11 (2015) e1005713. doi:10.1371/journal.pgen.1005713.

[18] B. Juarez, M.-H. Han, Diversity of Dopaminergic Neural Circuits in Response to Drug Exposure, Neuropsychopharmacology. 41 (2016) 1–23. doi:10.1038/npp.2016.32.

[19] P.K. Sonsalla, J.W. Gibb, G.R. Hanson, Roles of D1 and D2 dopamine receptor subtypes in mediating the methamphetamine-induced changes in monoamine systems., J. Pharmacol. Exp. Ther. 238 (1986) 932–7. http://www.ncbi.nlm.nih.gov/pubmed/2943891.

[20] C.J. Pike, D. Burdick, a J. Walencewicz, C.G. Glabe, C.W. Cotman, Neurodegeneration induced by beta-amyloid peptides in vitro: the role of peptide assembly state., J. Neurosci. 13 (1993) 1676–1687.

[21] B. Bryant, Camron D. and Yazdani, RNA binding proteins, neural development and the addictions, Genes Brain Behav. 15 (2016) 169–186. doi:10.1007/978-1-4614-5915-6.

[22] V. V Gusel, D.E. Korzhevskiy, NeuN As a Neuronal Nuclear Antigen and Neuron Differentiation Marker, Acta Naturae. 7 (2015) 42–47.

[23] M.C. Schaub, S.R. Lopez, M. Caputi, Members of the Heterogeneous Nuclear Ribonucleoprotein H Family Activate Splicing of an HIV-1 Splicing Substrate by Promoting Formation of ATP-dependent Spliceosomal Complexes, J. Biol. Chem. 282 (2007) 13617–13626. doi:10.1074/jbc.M700774200.

[24] D.M. Tiruchinapalli, M.D. Ehlers, J.D. Keene, Activity-dependent expression of RNA binding protein HuD and its association with mRNAs in neurons, RNA Biol. 5 (2008) 157–168. doi:10.4161/rna.5.3.6782.

[25] A. Aranburu, D. Liberg, B. Honorø, T. Leanderson, CArG box-binding factor-A interacts with multiple motifs in immunoglobulin promoters and has a regulated subcellular distribution, Eur J Immunol. 36 (2006) 2192–2202. doi:10.1002/eji.200535659.

[26] I. Grammatikakis, P. Zhang, A.C. Panda, J. Kim, S. Maudsley, K. Abdelmohsen, X. Yang, J.L. Martindale, O. Motiño, E.R. Hutchison, M.P. Mattson, M. Gorospe, Alternative Splicing of Neuronal Differentiation Factor TRF2 Regulated by HNRNPH1/H2, Cell Rep. 15 (2016) 1–9. doi:10.1016/j.celrep.2016.03.080.

[27] J.M. Kornhauser, C.W. Cowan, A.J. Shaywitz, R.E. Dolmetsch, E.C. Griffith, L.S. Hu, C. Haddad, Z. Xia, M.E. Greenberg, CREB transcriptional activity in neurons is regulated by multiple, calcium-specific phosphorylation events, Neuron. 34 (2002) 221–233. doi:10.1016/S0896-6273(02)00655-4.

[28] S. Paul, G.L. Snyder, H. Yokakura, M.R. Picciotto, A.C. Nairn, P.J. Lombroso, The Dopamine/D1 receptor mediates the phosphorylation and inactivation of the protein tyrosine phosphatase STEP via a PKA-dependent pathway., J Neurosci. 20 (2000) 5630–5638. doi:20/15/5630 [pii].

[29] N. Kim, J.O. Lee, H.J. Lee, S.K. Lee, J.W. Moon, S.J. Kim, S.H. Park, H.S. Kim, AMPK2 translocates into the nucleus and interacts with hnRNP H: Implications in metformin-mediated glucose uptake, Cell. Signal. 26 (2014) 1800–1806. doi:10.1016/j.cellsig.2014.03.023.

[30] H.-J. Kim, E.J. Song, K.-J. Lee, Proteomic analysis of protein phosphorylations in heat shock response and thermotolerance., J. Biol. Chem. 277 (2002) 23193–23207. doi:10.1074/jbc.M201007200.

[31] W. Cao, A. Razanau, D. Feng, V. Lobo, J. Xie, Control of alternative splicing by forskolin through hnRNP K during neuronal differentiation, Nucleic Acids Res. 40 (2012) 8059–8071. doi:10.1093/nar/gks504.

[32] F. Cobianchi, C. Calvio, M. Stoppini, M. Buvoli, S. Riva, Phosphorylation of human hnRNP protein A1 abrogates in vitro strand annealing activity, Nucleic Acids Res. 21 (1993) 949–955. doi:10.1093/nar/21.4.949.

[33] Y. He, R. Smith, Nuclear functions of heterogeneous nuclear ribonucleoproteins A/B, Cell. Mol. Life Sci. 66 (2009) 1239–1256. doi:10.1007/s00018-008-8532-1.

[34] G. Liu, A. Razanau, Y. Hai, J. Yu, M. Sohail, V.G. Lobo, J. Chu, S.K.P. Kung, J. Xie, A conserved serine of heterogeneous nuclear ribonucleoprotein L (hnRNP L) mediates depolarization-regulated alternative splicing of potassium channels, J. Biol. Chem. 287 (2012) 22709–22716. doi:10.1074/jbc.M112.357343.

[35] J. Xie, J.-A. Lee, T.L. Kress, K.L. Mowry, D.L. Black, Protein kinase A phosphorylation modulates transport of the polypyrimidine tract-binding protein., Proc. Natl. Acad. Sci. U. S. A. 100 (2003) 8776–8781. doi:10.1073/pnas.1432696100.

[36] J.M. Bain, M.T. Cho, A. Telegrafi, A. Wilson, S. Brooks, C. Botti, G. Gowans, L.A. Autullo, V. Krishnamurthy, M.C. Willing, T.L. Toler, B. Ben-Zev, O. Elpeleg, Y. Shen, K. Retterer, K.G. Monaghan, W.K. Chung, Variants in HNRNPH2 on the X Chromosome Are Associated with a Neurodevelopmental Disorder in Females., Am. J. Hum. Genet. 99 (2016) 1–7. doi:10.1016/j.ajhg.2016.06.028.

[37] B. Wolozin, Physiological Protein Aggregation Run Amuck: Stress Granules and the Genesis of Neurodegenerative Disease, Discov. Med. 17 (2014) 47–52. doi:10.1038/nbt.3121.ChIP-nexus.

[38] K. Marballi, N.K. Genabai, Y.A. Blednov, R.A. Harris, I. Ponomarev, Alcohol consumption induces global gene expression changes in VTA dopaminergic neurons, Genes, Brain Behav. 15 (2016) 318–326. doi:10.1111/gbb.12266.

